# A screen of covalent inhibitors in *Mycobacterium tuberculosis* identifies serine hydrolases involved in lipid metabolism as potential therapeutic targets

**DOI:** 10.1101/2021.06.07.447460

**Authors:** Brett M. Babin, Laura J. Keller, Yishay Pinto, Veronica L. Li, Andrew Eneim, Summer E. Vance, Stephanie M. Terrell, Ami Bhatt, Jonathan Z. Long, Matthew Bogyo

## Abstract

The increasing incidence of antibiotic-resistant *Mycobacterium tuberculosis* infections is a global health threat necessitating the development of new antibiotics. Serine hydrolases (SHs) are a promising class of targets because of their importance for the synthesis of the mycobacterial cell envelope. We screened a library of small molecules containing serine-reactive electrophiles and identified narrow spectrum inhibitors of *M. tuberculous* growth. Using these lead molecules, we performed competitive activity-based protein profiling and identified multiple SH targets, including enzymes with uncharacterized functions. Lipidomic analyses of compound-treated cultures revealed an accumulation of free lipids and a substantial decrease in lipooligosaccharides, linking SH inhibition to defects in cell envelope biogenesis. Mutant analysis revealed a path to resistance via the synthesis of mycocerates, but not through mutations to target enzymes. Our results suggest that simultaneous inhibition of multiple SH enzymes is likely to be an effective therapeutic strategy for the treatment of *M. tuberculosis* infections.

## Introduction

Tuberculosis is a top-ten cause of death worldwide and the leading cause of death by a bacterial pathogen. The successful treatment of tuberculosis is a challenging, multifaceted problem whose solution will require improvements in diagnostics, antibiotic access, and treatment monitoring. The increasing incidence of antibiotic resistance in the causative agent of tuberculosis, *Mycobacterium tuberculosis,* is a growing global health threat. Identification of new drug targets and new antibiotics that circumvent resistance is an important step in combatting tuberculosis.

Serine hydrolases (SHs) are a promising class of potential therapeutic enzyme targets for new antibiotics. SHs are involved in a variety of critical cellular processes, including lipid homeostasis, metabolism of host lipids, and cell wall synthesis and maintenance (Bachovchin and Cravatt, 2012). Because SHs have many roles in assembly, modification, and maintenance of the extensive mycobacterial cell envelope, they are overrepresented in the *M. tuberculosis* genome compared to other pathogenic bacteria (Cotes et al., 2008). They comprise 4.0% of the proteome (158 of 4,080 total proteins) compared to 2.0%, 1.0%, and 1.7% for the other common bacterial pathogens *Pseudomonas aeruginosa, Salmonella enterica,* and *Staphylococcus aureus* (Lentz et al., 2018). SHs are also significantly more prevalent in *M. tuberculosis* than in the human proteome (<1%) (Simon and Cravatt, 2010). Furthermore, many *M. tuberculosis* SHs lack human homologs, suggesting it is feasible to develop inhibitors that are unreactive toward host enzymes. In addition, small molecules that target human SHs have been developed to treat a range of conditions including hypertension, obesity, diabetes and others (Bachovchin and Cravatt, 2012). One of the most successful therapeutic small molecule SH inhibitors is tetrahydrolipstatin (THL, marketed as Orlistat) which is used to treat obesity by blocking key lipid mobilizing enzymes in the gut, thus blocking the utilization of dietary fats (Borgstrom, 1988; Drent and van der Veen, 1993). Furthermore, the extensive efforts to develop SH inhibitors as therapeutics has created diverse compound classes that inhibit these enzymes and has established successful strategies to optimize compound selectivity (Adibekian et al., 2011; Bachovchin et al., 2010; Chang et al., 2013; Faucher et al., 2020). Therefore, SHs are a highly diverse and ‘druggable’ family of targets whose function in bacteria remains poorly defined.

Previous studies have identified small molecule inhibitors of *M. tuberculosis* SHs, including THL (Low et al., 2009), the mammalian liposomal acid lipase inhibitor lalistat (Lehmann et al., 2016), the cyclipostin CyC_17_ (Nguyen et al., 2017), the human hormone sensitive lipase inhibitor M*m*PPOX (Delorme et al., 2012), the β-lactone EZ120 (Lehmann et al., 2018), and the triazole ureas AA691 and AA692 (Li et al., 2021). These inhibitors all make use of an electrophile which covalently modifies the active site serine present in all members of this enzyme class. While these studies demonstrate the importance of SHs as therapeutic targets, they generally have been unable to determine the specific mechanisms by which SH inhibition interferes with *M. tuberculosis* growth or viability. A key benefit of using inhibitors that covalently modify the active site of their protein targets is that those targets can be readily identified by activity-based protein profiling (ABPP) mass spectrometry experiments. ABPP experiments have been used to identify targets for each of the SH inhibitors listed (Lehmann et al., 2018; Lehmann et al., 2016; Li et al., 2021; Ravindran et al., 2014). However, all of the reported SH inhibitors with activity in *M. tuberculosis* target multiple SHs, making it difficult to identify the enzyme or set of enzymes whose functions are essential for bacterial survival. In most cases, the genes that code for SHs are not essential (Griffin et al., 2011; Rengarajan et al., 2005; Sassetti et al., 2003; Sassetti and Rubin, 2003), suggesting that efficacy of bactericidal or bacteriostatic SH inhibitors likely relates to their ability to simultaneously inhibit multiple enzymes.

For these reasons, we sought to perform a phenotypic, cell-based screen of a highly focused library of potential covalent SH inhibitors to identify compounds that inhibit *M. tuberculosis* growth. We reasoned that such a screen would yield hits that could then be used to identify specific SH targets whose function is essential for survival of the bacteria. Because the library consists exclusively of covalent binding electrophiles, it was also possible to use hit compounds to directly identify the relevant SHs whose inhibition is linked to the inhibition of bacterial growth. From this screen, we identified a series of 7-urea chloroisocoumarin inhibitors with narrow spectrum activity towards *M. tuberculosis* that lacked host cell cytotoxicity. We performed competitive activity-based protein profiling proteomic experiments and identified multiple protein targets covalently modified by this class of compounds, including uncharacterized lipases. We used LC-MS-based lipidomics and whole-genome sequencing of a resistant mutant to demonstrate that these SHs are involved in aspects of cell wall biogenesis that are disrupted by inhibitor treatment, thus making them promising therapeutic targets.

## Results

### HTS yields electrophilic inhibitors of M. tuberculosis growth

We screened a library of 942 compounds featuring cysteine- and serine-reactive electrophiles (Child et al., 2013; Hall et al., 2011) for their ability to inhibit exponential growth of *M. tuberculosis* in culture (Fig. 1a). We tested compounds at a single dose (20 µM) with a one-week incubation, which yielded a 4.9% hit rate (46/942 compounds). To validate the 46 hit compounds, we performed a 10-point dose response analysis to estimate IC_50_ values (Fig. 1b, Dataset S1). Of the top 14 compounds with EC_50_ < 5 µM, seven were excluded due to host cell cytotoxicity as reported previously for human pancreatic cell lines (Gruner et al., 2016). The validated hits consisted of two core chemical scaffolds: 7-urea chloroisocoumarins and peptide diphenylphosphonates (Table 1, Fig. 1c), both of which are expected to target the active site of serine hydrolases or proteases. Though the screening library contains many chloroisocoumarin compounds with diverse substitutions at the 7 position, we note that only those featuring a urea arose as hits.

**Figure 1.**
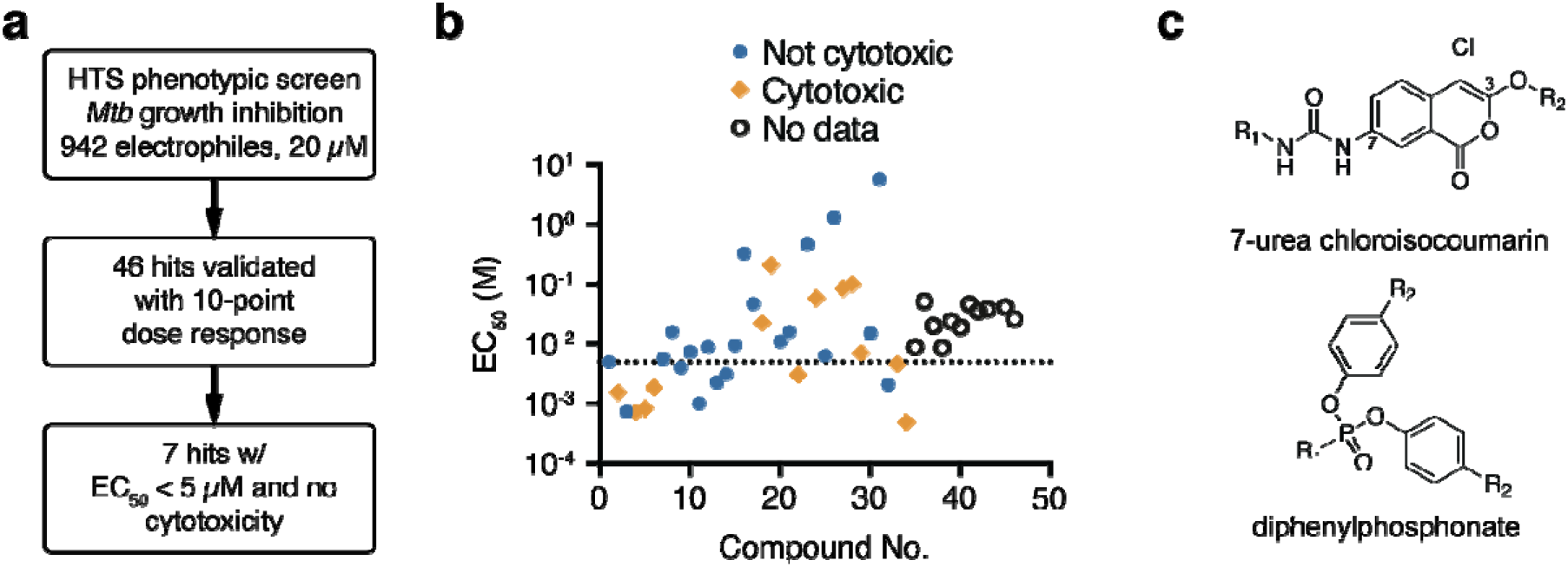
Screen of electrophilic compounds yields inhibitors of *M. tuberculosis* growth. (a) Screening strategy. (b) EC_50_ values for 46 hits. Cytotoxicity data were obtained from (Gruner et al., 2016). The dotted line indicates the 5 µM cutoff used to prioritize hits. (c) Core scaffolds of the top seven hits.

**Table 1.**
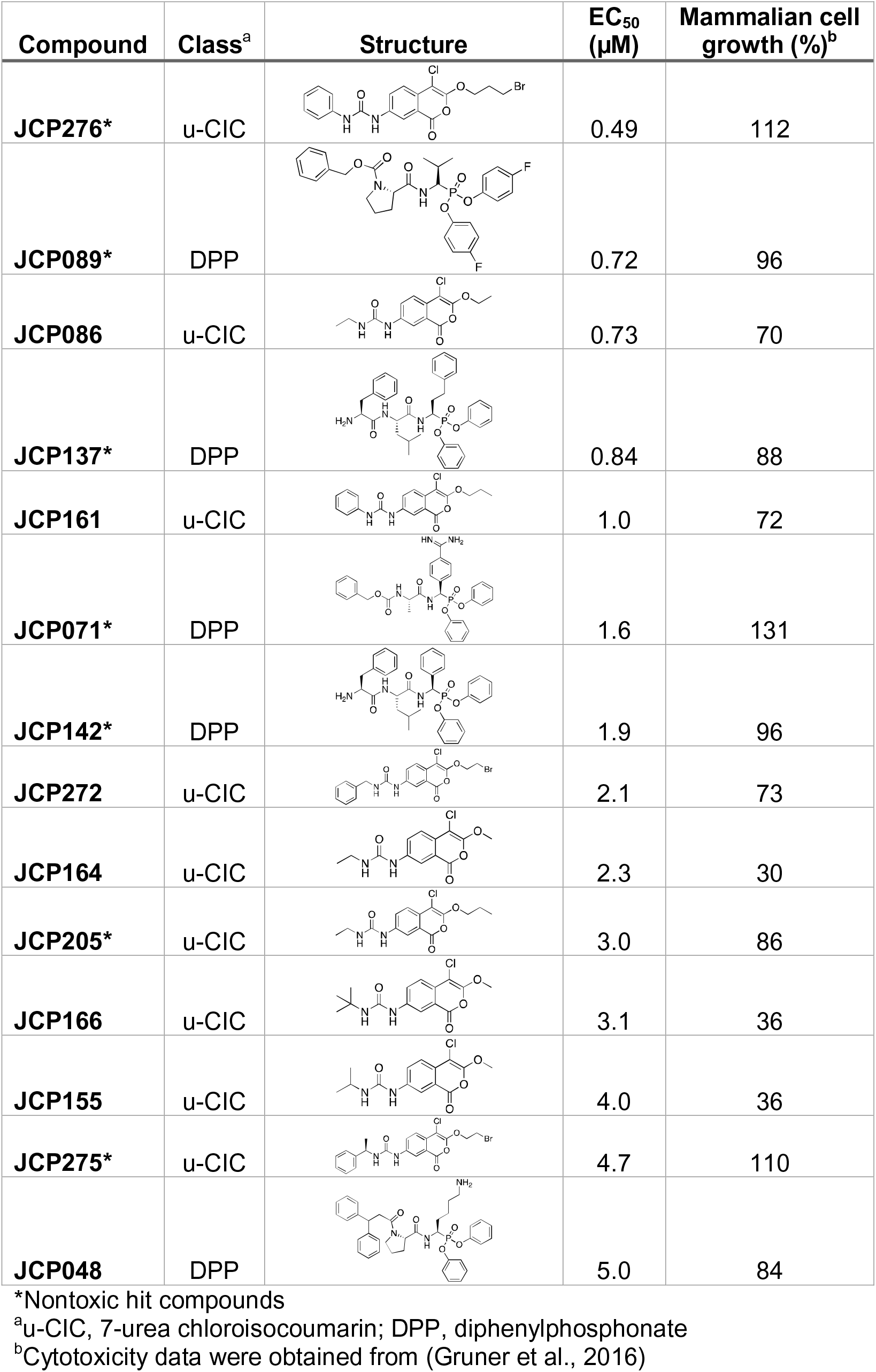
Screening hits.

| Compound | Class<sup>a</sup> | Structure | EC<sub>50</sub><br>(μM) | Mammalian cell<br>growth (%)<sup>b</sup> |
| --- | --- | --- | --- | --- |
| JCP276* | u-CIC |  | 0.49 | 112 |
| JCP089* | DPP |  | 0.72 | 96 |
| JCP086 | u-CIC |  | 0.73 | 70 |
| JCP137* | DPP |  | 0.84 | 88 |
| JCP161 | u-CIC |  | 1.0 | 72 |
| JCP071* | DPP |  | 1.6 | 131 |
| JCP142* | DPP |  | 1.9 | 96 |
| JCP272 | u-CIC |  | 2.1 | 73 |
| JCP164 | u-CIC |  | 2.3 | 30 |
| JCP205* | u-CIC |  | 3.0 | 86 |
| JCP166 | u-CIC |  | 3.1 | 36 |
| JCP155 | u-CIC |  | 4.0 | 36 |
| JCP275* | u-CIC |  | 4.7 | 110 |
| JCP048 | DPP |  | 5.0 | 84 |

\*Nontoxic hit compounds
<sup>a</sup>u-CIC, 7-urea chloroisocoumarin; DPP, diphenylphosphonate
<sup>b</sup>Cytotoxicity data were obtained from (Gruner et al., 2016)

We chose to focus all subsequent efforts on the most potent compound from the validated hits, JCP276 (Fig. 2a). Upon resynthesis, it showed similar potency for growth inhibition after one week of incubation as observed in the original screen, with an EC_50_ of 1.7 µM and EC_90_ of 10 µM (Fig. 2b). Three-week incubations revealed a MIC of 50 µM (23 mg/L). Treating liquid cultures with a sub-MIC of JCP276 resulted in growth arrest for approximately 14 days, followed by outgrowth (Fig. 2c), suggesting that the activity is bacteriostatic, and that differences in one-week and three-week potencies may be due to compound degradation. Indeed, growth inhibition could be maintained by a second addition of JCP276 at day 14, suggesting that this effect was a result of compound inactivation or degradation rather than evolved resistance.

**Figure 2.**
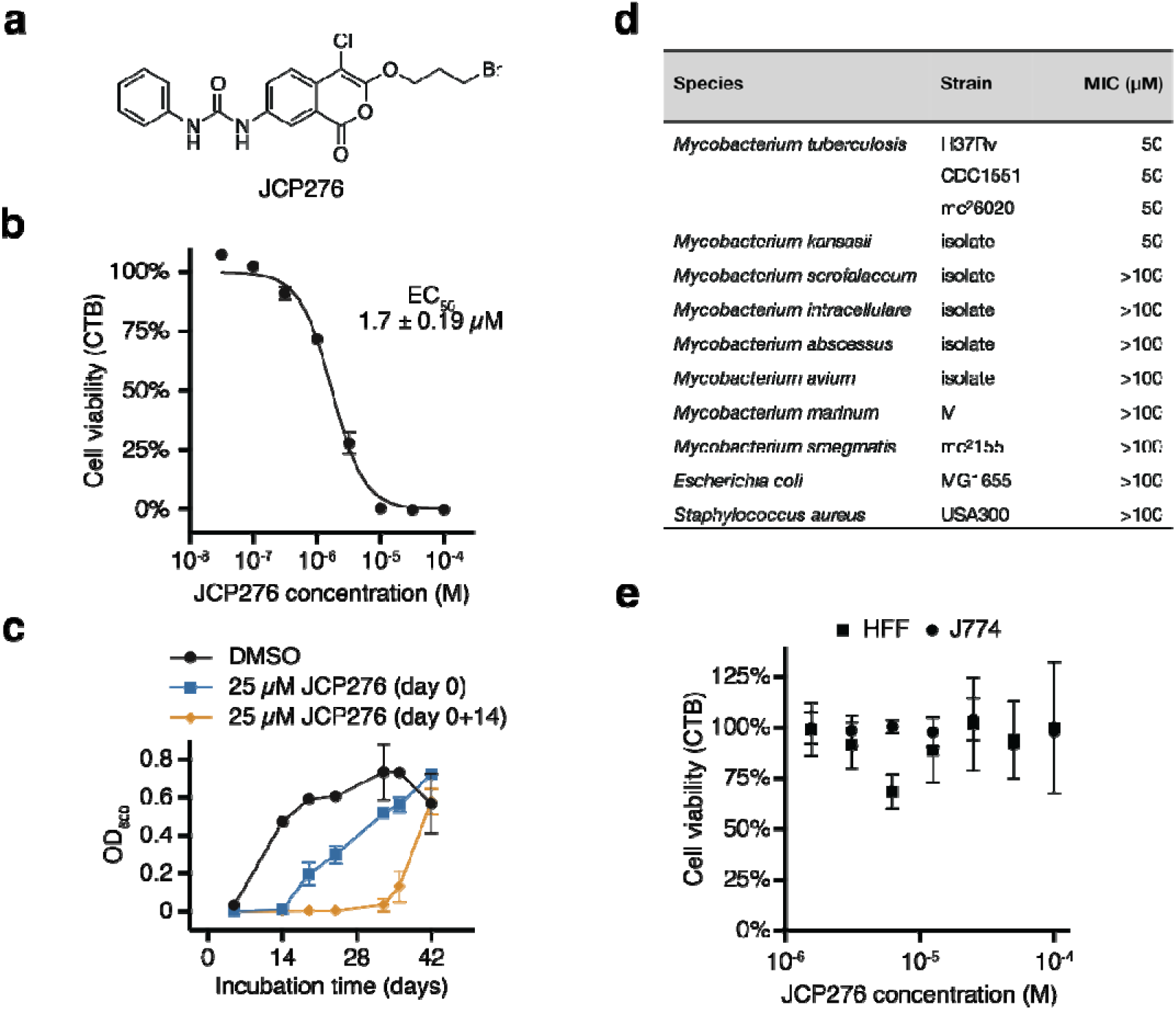
JCP276 is a narrow spectrum inhibitor of *M. tuberculosis* growth. (a) Structure of JCP276. (b) Dose-dependent inhibition of *M. tuberculosis* growth by JCP276 following a 7-day incubation. Data represent the mean ± standard deviation of three biological replicates. Data were fit to two-parameter parametric model (solid line) and the EC_50_ is reported as the mean ± 95% confidence interval. (c) Growth of *M. tuberculosis* as measured by OD_600_. JCP276 was added at day 0 of the experiment (blue squares) or at day 0 and again at day 14 (orange diamonds). Data represent the mean ± standard deviation of three biological replicates. (d) MIC_90_ of JCP276 for various bacterial strains as measured by visual inspection of growth in well-plates. N.D.: no growth inhibition observed for JCP276 doses up to 200 µM. (e) Cytotoxicity measurements for mammalian cell lines in the presence of JCP276. Data represent the mean ± standard deviation of three biological replicates.

To evaluate the selectivity of JCP276, we tested its effect on other bacteria as well as mammalian cells. While JCP276 inhibited growth of three laboratory strains of *M. tuberculosis* and a clinical isolate of *M. kansasii,* the compound had no effect on the growth of *Escherichia coli* or *Staphylococcus aureus* at doses up to 100 µM (Fig 2d). Faster growing Mycobacteria and pathogenic isolates *M. smegmatis*, *M. marinum*, *M. intracellulare, M. abscessus, M. avium* were similarly unaffected, suggesting a very narrow spectrum of activity. JCP276 was also not cytotoxic to a murine macrophage cell line or human foreskin fibroblasts up to 100 µM after 24 h of incubation (Fig. 2e).

### Activity-based protein profiling enables the identification of hydrolase targets of JCP276

Because the chloroisocoumarin electrophile is expected to covalently modify the active site of serine hydrolases, we used competitive ABPP to identify protein targets of JCP276 and the structurally similar hit JCP275 (Fig. S1a). We used fluorophosphonate (FP) coupled to either biotin or TMR (Fig. 3a) as a general probe for serine hydrolases. Cells were treated with JCP276 or JCP275 for 1 h and lysates were treated with FP-tetramethylrhodamine (TMR) to label the active site of serine hydrolases. Labeled proteins were visualized via SDS-PAGE (Fig. 3b, Fig. S1b). Five proteins were competed by the inhibitors at doses at or above the EC_50_ values of the two inhibitors, suggesting that both target multiple SH proteins.

**Figure 3.**
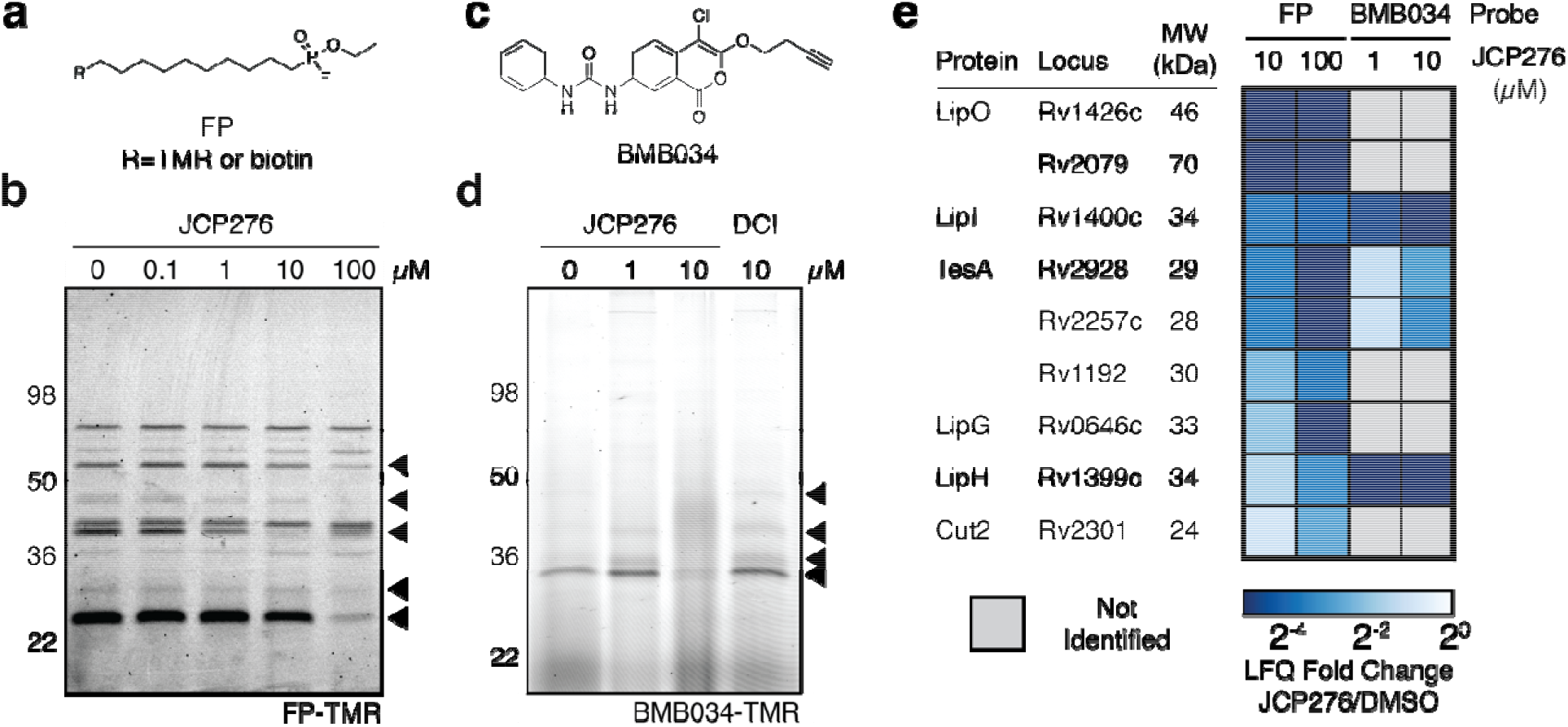
Activity-based protein profiling reveals serine hydrolase targets of JCP276. Structures of (a) FP and (c) BMB034 probes. (b) Competitive ABPP using FP-TMR. Bacterial cultures were treated with DMSO or JCP276 as indicated for 1 h at 37 °C. Lysates were treated with 1 µM FP-TMR for 30 min at 37 °C and proteins were separated by SDS-PAGE. (d) Competitive ABPP using BMB034. Bacterial cultures were pre-treated with DMSO, JCP276, or DCI as indicated for 1 h at 37 °C and then treated with 1 µM BMB034 for 1 h at 37 °C. Lysates were subjected to click chemistry conditions with azide-TMR and proteins were separated by SDS-PAGE. Gels show TMR fluorescence. (e) Summary of proteomic results from FP and BMB034 ABPP experiments. The heatmap represents the ratio of the abundance of each protein between cultures treated with JCP276 at the indicated dose and cultures treated with DMSO.

While most serine hydrolases are targeted by FP (Bachovchin et al., 2010), we also considered the possibility that the chloroisocoumarin electrophile inhibits proteins that are not labeled in the FP competition experiment. To account for this possibility, we synthesized alkyne probe versions of JCP276, featuring the alkyne attached either to the 3 position (BMB034, Fig. 3c) or the 7 position (BMB038, Fig. S1c) of the chloroisocoumarin core. Both BMB034 and BMB038 inhibited *M. tuberculosis* growth in culture, albeit with decreased potencies compared to the parent compound JCP276 with EC_50_s of 8.7 and 36 µM, respectively (Fig. S2b). We used BMB034 for competitive ABPP because it was the more potent of the two probes. After incubation of live cells in culture with BMB034, we lysed the cells, and performed CLICK chemistry with azide-TMR to visualize BMB034-labeled proteins. Four proteins were consistently labeled by 1 µM BMB034 (Fig. 3d), while higher doses (10 and 100 µM) labeled a larger set of proteins (Fig. S1e). Pre-treatment of cultures with JCP276 blocked the labeling of four proteins by BMB034 (Fig. 3d, Fig. S1f). As a negative control we pre-treated cells with dichloroisocoumarin (DCI), a general inhibitor of serine hydrolases and proteases that does not inhibit *M. tuberculosis* growth, and found that it did not compete for labeling of any of the proteins labeled by BMB034. These data suggest that the BMB034-labeled proteins are likely phenotypically relevant targets of JCP276.

To identify putative JCP276 targets, we used both the general SH probe FP-biotin or BMB034 and then performed click labeling with desthiobiotin to affinity purify the labeled proteins for mass spectrometry analysis. For both probes, we pretreated cells with the lead inhibitor JCP276 and then measured the change in recovery of the target SH proteins. We found that the set of FP-biotin labeled proteins overlapped substantially with the list of FP-reactive proteins reported in three previous studies, confirming that our probe labeling was working as expected (Li et al., 2021; Ortega et al., 2016; Patel et al., 2018; Tallman et al., 2016) (Table S1). Overall, we identified 25 of the 27 proteins (93%) found in all three existing datasets and 44 of the 47 proteins (94%) found in at least two existing datasets. These comparisons suggest that despite differences in probe design, enrichment protocols, and culture conditions, our FP enrichment experiment captured the majority FP-reactive enzymes. Because protein targets of JCP276 are expected to be depleted in the pre-treatment conditions, we calculated abundance ratios between proteins enriched from cells that had been pre-treated with 10 µM or 100 µM of JCP276 or JCP275, relative to the DMSO treated control (Table S1). Abundance ratios for each predicted serine hydrolase (Tallman et al., 2016) correlated well between both the JCP275 and JCP276 pretreated samples (n=55, r=0.87, Fig. S2a), suggesting that each compound inhibits the same set of protein targets with similar potencies and likely acts via the same mechanism. Nine of the 55 identified proteins were significantly reduced in abundance by 10 µM pre-treatment with JCP275 and JCP276 (Fig. 2e, Table 2, Fig. S2b). The abundance ratios were the same or lower following pre-treatment with the 100 µM dose of each compound, confirming the expected dose response of true targets.

Enrichment via BMB034 yielded a smaller set of predicted serine hydrolases. While the entire enrichment experiment yielded 106 protein identifications, only seven were competed by pre-treatment with 10 µM JCP276, and only four of those were not competed by DCI (Table S1). These four proteins are all predicted serine hydrolases that were identified as putative targets in the FP enrichment experiment (Fig. 2e). Five of the proteins competed in the FP enrichment experiment were not detected in the BMB034 enrichment experiment, suggesting potential differences in the target binding properties of the alkyne probe BMB034 compared to the original JCP276 lead molecule. The concentration of BMB034 used in direct labeling experiments (1 µM) was below its EC_50_ so it is possible that some target proteins were not sufficiently labeled at this lower dose and therefore where not identified in the direct labeling study. A large number of the identified proteins were not competed by JCP276 and are not annotated as serine hydrolases, suggesting that they are likely artifacts of the enrichment procedure.

Taken together, the two enrichment experiments define a set of nine putative protein targets that includes predicted esterases and lipases (e.g., LipO, LipI, LipG, NlhH/LipH), though neither the biochemical activity nor the biological function of most of these putative targets have been confirmed. Importantly, none of the genes encoding the putative targets are annotated as essential according to transposon sequencing datasets (Griffin et al., 2011; Rengarajan et al., 2005; Sassetti et al., 2003; Sassetti and Rubin, 2003). The fact that inhibition of each target individually is not expected to cause a growth defect suggests that JCP276 acts by simultaneous inhibition of multiple SH targets.

### Lipidomic analysis links THL and chloroisocoumarin mechanisms of growth inhibition

It is currently unclear how polypharmacological inhibition of SHs disrupts bacterial growth. Because many of the protein targets of JCP276 and other SH inhibitors are biochemically and biologically uncharacterized, target identification alone is insufficient to develop a model for a mechanism of action. Because many of the putative targets are expected to be involved in the synthesis and modification of lipids, we hypothesized that inhibitor treatment might alter cellular lipid profiles. To compare the modes of action between the chloroisocoumarin inhibitors and a previously reported lipase inhibitor, we compared lipid profiles from cells treated with JCP276, BMB034, or THL at twice their MICs. We extracted lipids from bacterial cultures and performed an analysis using LC-MS. The resulting spectra were annotated by matching ions to the MycoMass database (Layre et al., 2011) to identify validated, mycobacterium-specific lipids. Fold changes for each identified lipid were calculated as the ratio of mean integrated ion currents from treated and untreated samples. Across all four conditions, we identified and matched 1,206 lipids to the database. Principal component analysis (PCA) of individual samples resulted in two clusters: one containing the DMSO-treated control samples, and another containing all of the compound-treated samples (Fig. 4a). In addition, treated and untreated samples are differentiated by the first principal component, which explains 35% of the variance among the samples. Results from this analysis suggest that the lipidomic data are consistent among replicates because each treatment condition is separated by PC1 and PC2. It also confirms that bacteria treated with either JCP276, BMB034, or THL exhibit a shift in lipid profiles. Finally, it shows that treatment with any of the three compounds results in a similar perturbation to the lipid profiles. The latter observation supports the hypothesis that both chloroisocoumarins and THL act via a similar general mechanism.

**Figure 4.**
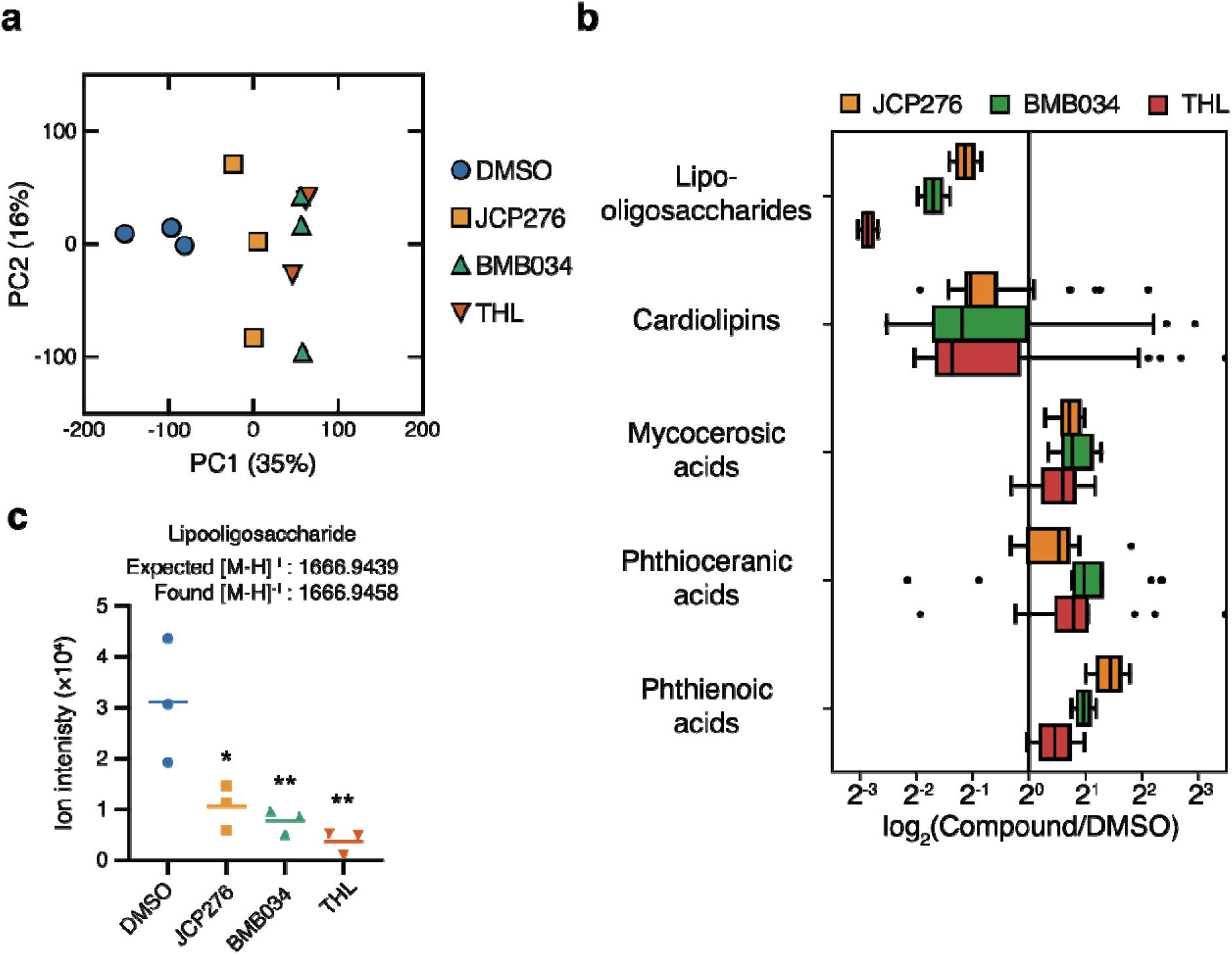
Serine hydrolase inhibitors alter cellular lipid profiles. (a) PCA of lipid abundances from three biological replicates for each compound treatment. Solid cultures were treated with DMSO or 100 µM of JCP276, BMB034, or THL for 5 d at 37 °C. Lipids were extracted and analyzed by LC-MS. (b) Boxplots of lipid abundance ratios between cultures treated with each compound or cultures treated with DMSO. The vertical line represents the median value for each lipid class, boxes represent the upper and lower quartiles, whiskers represent 1.5 times the interquartile range, and circles represent outliers from this range. (c) Ion intensities for lipooligosaccharide for each treatment condition. Data from treated cultures were compared to the DMSO control by ANOVA followed by Dunnett’s test for multiple comparisons.

To more carefully define the changes in lipid components upon inhibition of SHs, we calculated the mean fold change in ion intensities of compound-treated samples compared to the DMSO-treated controls (Table S2). A global analysis of changes showed strong correlations between JCP276 vs. BMB034 (r=0.90) and JCP276 vs. THL (r=0.89) (Fig. S3a-b), further supporting the hypothesis that THL and chloroisocoumarin compounds act via similar lipid perturbations. Although fold changes for individual lipid species varied, the majority of classes had median fold changes that were close to 1 (Fig. S3c), suggesting that perturbations were limited to a small subset of lipids. The lipid classes that showed the greatest changes were lipooligosaccharides and cardiolipins which were depleted in compound-treated cells and the free mycobacterial lipids mycocerosic, phthioceranic, and phthienoic acids which were enriched (Fig. 4b).

### PDIM regulates susceptibility to JCP276

We attempted to generate spontaneous resistance mutants by culturing a high concentration of bacteria on solid agar containing supra-MICs of JCP276. We performed initial attempts using 4×10^6^ CFU with 100 µM JCP276 but these attempts did not yield any resistant colonies (two independent experiments). However, we were able to isolate resistant clones by increasing the number of cells applied to each plate to 2×10^9^ CFU (500-fold more cells). Resistant strains were isolated from three different solid cultures (BBMT04, BBMT05, and BBMT06). We treated liquid cultures of each strain with JCP276 and confirmed that all three exhibited a shift in EC_50_ values in excess of 100 fold (Fig. 5a, EC_50_ > 300 µM). Whole genome sequencing of the wild-type strain revealed 93 mutations present in our laboratory strain compared to the H37Rv reference genome, including frame shifts and point mutations that resulted in the truncation of seven proteins (Table S3). Many of the mutations (42%) were found in *pe/ppe* genes, known hotspots for sequence variation in the *M. tuberculosis* genome (McEvoy et al., 2012). We chose one mutant, BBMT05, for whole genome sequencing and found 92 of the 93 mutations identified in the wild-type strain. The one exception was a 7 bp insertion in the *ppsA* gene. Unexpectedly, the insertion yielded a gene sequence that exactly matched the reference genome. Because this seemed like a strange finding as it resulted in a gain of function back to the original reference genome, we performed PCR and Sanger sequencing to confirm the mutation and to verify the sequences of this region for the BBMT04 and BBMT06 strains as well. For all three mutants that exhibited resistance to JCP276, we found that the *ppsA* truncation was reverted back to the reference sequence (Fig. 5b). This observation suggests that our original ‘wild-type’ strain was not isogenic but rather contained a majority of cells that contain the PpsA truncation mutation combined with a small subpopulation coding for the full-length gene. This subpopulation with a fully functional *ppsA* gene is less sensitive to JCP276 and therefore became highly enriched in our resistance study. If the *ppsA-*intact subpopulation is present at very low levels, this would also explain our inability to generate resistant mutants when starting with fewer cells in the inoculum.

**Figure 5.**
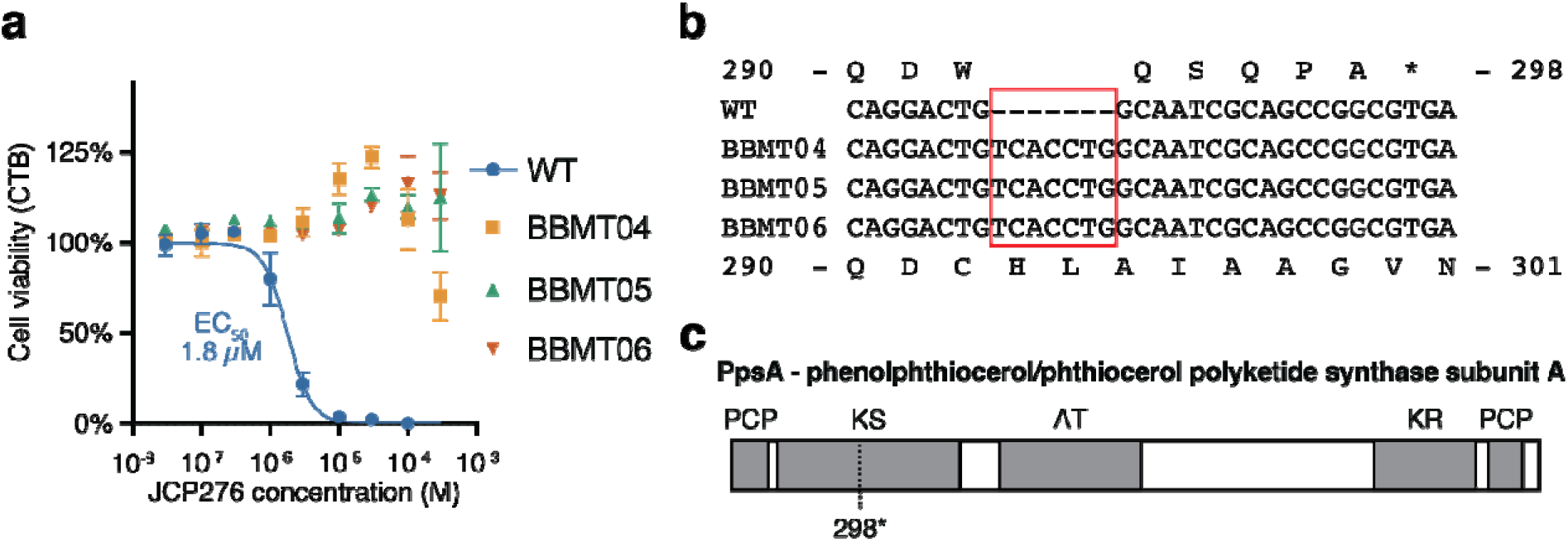
PDIM competency confers resistance to JCP276. (a) Dose-dependent inhibition of wild-type and mutant *M. tuberculosis* growth by JCP276 following a 7-day incubation. Data represent the mean ± standard deviation of three biological replicates. Data were fit to a two-parameter parametric model (solid line). (b) Sanger sequencing results for the *ppsA* gene. Amino acid sequences are annotated above (wild-type strain) and below (mutant strains). Amino acids are indicated above or below the first base of each triplet codon. (c) Diagram of PpsA protein with the location of the premature stop codon at residue 298 indicated. Domains are indicated: PCP, peptidyl carrier protein; KS, β-ketoacyl ACP synthase; AT, acyl transferase; and KR, ketoreductase.

PpsA is a phenolpthiocerol ketide synthase that serves as part of the PpsABCDE complex required for the synthesis of PDIM (Trivedi et al., 2005). PpsA catalyzes the first steps in the extension and modification of lipids to convert long chain fatty acids (mycosanoic acids) into mycocerosic acids and phthioceranic acids which are assembled to make PDIM. The truncated copy of PpsA encoded by the wild-type strain lacks the catalytic domains required for lipid processing and for transfer to the next polyketide synthase in the assembly line of PDIM synthesis (Fig. 5c). Thus, the wild-type strain lacks the ability to produce PDIM. Laboratory culture is known to select for *M. tuberculosis* strains that lack PDIM, often via mutation of the *pps* genes, because PDIM deficient cells have a growth advantage in standard medium (Domenech and Reed, 2009). In addition, PDIM serves as a formidable barrier for small molecule uptake and it is well characterized that PDIM-deficient strains are more susceptible to an array of antibiotics (Chavadi et al., 2011; Mohandas et al., 2016). Therefore, we hypothesize that cells deficient in PDIM synthesis readily uptake JCP276, allowing inhibition of growth so that only a small subpopulation of PDIM competent cells can be cultured in the presence of supra-MIC of JCP276.

### Diverse SH inhibitors target multiple enzymes to disrupt bacterial growth

Given that none of the identified protein targets are encoded by essential genes, we hypothesize that JCP276 acts by targeting multiple SHs whose combined inhibition prevents bacterial growth. This hypothesis is supported by the fact that the previously reported SH inhibitors with anti-mycobacterial activity also target multiple serine hydrolases. We sought to compare the protein targets of this set of compounds to distinguish between targets expected to be important for the cellular effects and targets whose inhibition is not necessary for growth inhibition. We compared the targets of JCP276, THL, lalistat, CyC_17_, AA692, and EZ120 (Fig. 6a) that have been identified in this study and in prior ABPP experiments. Of the 77 predicted SHs in *M. tuberculosis* (Tallman et al., 2016), 40 were identified as targets of at least one of the six compounds. We note that variations in the design of the ABPP experiments, the probes used, bacterial growth conditions, and LC-MS/MS instruments may result in *bona fide* targets being missed in any single study. In addition, some protein targets may be more susceptible to small molecule electrophiles (e.g., those with broad substrate promiscuity, high expression levels, or especially reactive nucleophile active sites). Because Li, et al. included a structurally similar but less potent compound in their ABPP study, their set of prioritized AA692 targets excludes SHs that are readily targeted by serine-reactive electrophiles but that do not contribute to growth inhibition. Therefore, to narrow down this list of possible targets we sought proteins that are: (i) targeted by at least three of the compounds, (ii) targeted by JCP276, (iii) targeted by AA692, and (iv) not targeted by the less potent triazole urea, AA702 (Li et al., 2021). Of the nine proteins targeted by more than three compounds, only TesA meets these criteria (Fig. 6b). LipM, LipN, and Rv1730c are potentially important targets, but we can exclude their role in the activity of JCP276 because they were identified but not competed in our FP ABPP experiment (Table S1). Because TesA is nonessential for growth in culture, it unlikely to be the only target by which JCP276 and the other SH inhibitors exert their effects on mycobacterial growth. However, the above analysis suggests that TesA is an important and shared target of most SH inhibitors, and that growth inhibition may require targeting of TesA along with other lipases.

**Figure 6.**
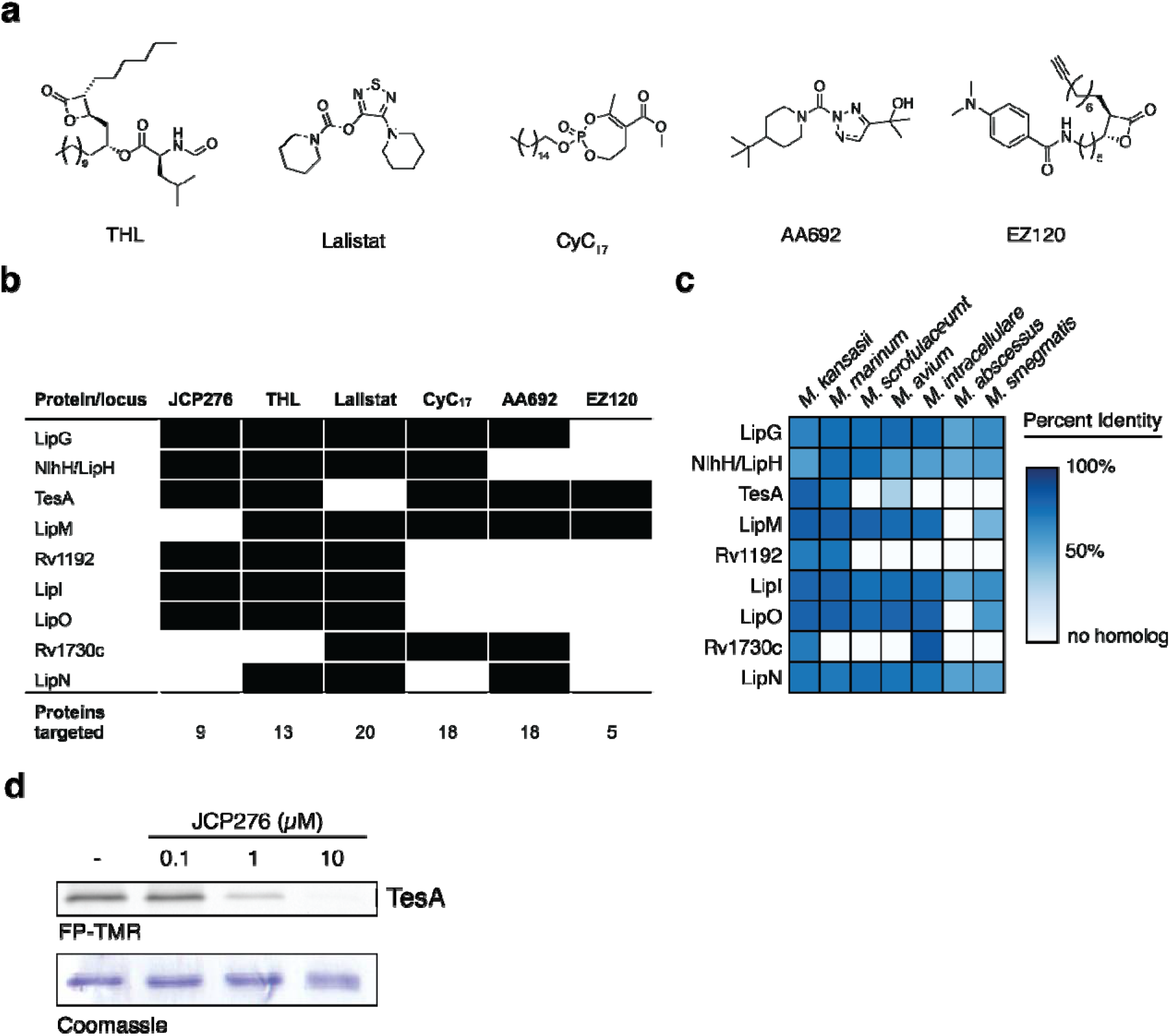
SH inhibitors target a conserved set of enzymes. (a) Structures of SH inhibitors with activity against Mycobacteria. (b) Proteins targeted by at least three of the SH inhibitors from (a) as determined by ABPP. Black boxes indicate that a protein was identified as a target. (c) Homologs of each protein from (b) in various NTMs. The heatmap indicates the percent identity of each protein to its *M. tuberculosis* homolog. White boxes indicate no homolog in that species. (d) Competitive ABP of *M. tuberculosis* TesA. Recombinant TesA was incubated with DMSO or JCP276 as indicated, treated with FP-TMR, and separated by SDS-PAGE. Coomassie stain indicates equal protein loading.

Given that JCP276 has narrow spectrum activity against *M. tuberculosis* and *M. kansasii*, we hypothesized that conservation of putative targets among mycobacteria may also explain the differences in susceptibility among bacteria. We performed BLAST analysis of all 77 predicted SHs in *M. tuberculosis* to search for homologs in *M. kansasii* and the non-tubercular mycobacteria whose growth is not inhibited by JCP276. A majority of SHs (73%) from *M. tuberculosis* had homologs in all other mycobacteria. Of the targets of JCP276, TesA and Rv1192 are notable because their homologs are present in *M. kansasii* but few other mycobacteria (Fig. 6c). Analysis of both the composition of shared SH inhibitor targets and homology across species indicate TesA as a protein that is likely to be important to the inhibitory action of JCP276. To verify that TesA is indeed inhibited by JCP276, we purified recombinant TesA, pre-treated the enzyme with JCP276, and labeled the active site with FP-TMR (Fig. 6d). We observed a dose-dependent decrease in FP labeling consistent with covalent modification of the active site by JCP276, confirming TesA as a target. We also note that overexpression of TesA in *M. bovis* reduced the susceptibility of that organism to THL (Ravindran et al., 2014). Given that treatment with JCP276 and BMB034 induce similar lipidome remodeling as THL, it is likely that the mechanisms of action of each compound are related.

## Discussion

This study and others reporting the use of SH inhibitors to disrupt *M. tuberculosis* growth highlight the benefits of a polypharmacological strategy for tuberculosis therapy. In each study, phenotypic screens yielded inhibitors that target multiple enzymes which, by themselves, are nonessential for growth in culture. Because each target is nonessential, a target-based approach would not have progressed to the development of SH inhibitors. One of the key benefits of antimicrobial compounds that inhibit multiple targets is that there appears to be a larger barrier to evolving resistance. No resistance was observed for the triazole urea SH inhibitors AA691 and AA692 with resistance rates expected to be lower than 1×10^9^ (Li et al., 2021). In our study, resistance mutants were obtained at a rate ∼2×10^9^, but we note that the mechanism of resistance is likely related to the increased barrier of entry in the mutant strains due to their PDIM production. The fact that PDIM prevents growth inhibition by JCP276 means that this compound is not likely to be a viable antibiotic, given that PDIM is critical for infection (Cox et al., 1999) and clinical isolates of *M. tuberculosis* are PDIM competent (Domenech and Reed, 2009). However, it is important to note that resistance was not observed via mutation of any of the protein targets of the compound, suggesting that an optimized compound with enhanced cell penetrance would likely not readily yield resistant mutants.

Determining the mechanism of action by which an inhibitor with multiple targets impacts fitness is challenging. Targets whose inhibition is important for growth inhibition can, in principle, be distinguished from other targets by combinatorial genetic knockdown or deletion experiments. However, in the case of SH inhibitors with ∼10-20 targets, a full genetic analysis with all combinations is prohibitive. In this study, we used a comparative lipidomic approach in an effort to identify the pathways that are perturbed by JCP276 treatment. Linking the lipidomic results to the inhibition of enzyme activities remains challenging because many of the protein targets in *M. tuberculosis* are biochemically and physiologically uncharacterized. However, the substantial lipid changes upon treatment with SH inhibitors point to a plausible model mechanism in which inhibition of thioesterases lead to the buildup of fatty acids and prevent the synthesis of critical membrane components. It also identifies a signature of lipid accumulation that correlates with bactericidal activity, suggesting that this accumulation could be used as a readout in future drug screens to enable the identification of compounds with multi-target modes of action

We observe that SH inhibitor treatment results in an accumulation of mycobacterial fatty acids including mycocerosic, phthioceranic, and phthienoic acids (Fig. 4b). *M. tuberculosis* uses a suite of polyketide synthases (PKSs) for the biosynthesis of these complex mycobacterial lipids (Minnikin et al., 2002; Natarajan et al., 2008). A key step in the synthesis of these lipids is the cleavage of the thioester bond which links a given fatty acid to a cysteine residue on its cognate PKS. For example, PDIM synthesis relies on the thioesterase activity of TesA to release lipids produced by the PKSs PpsABCDE and Mas (Nguyen et al., 2018). In the absence of active thioesterases, fatty acids cannot be removed from PKSs and are therefore not substrates for conjugation to the diverse lipid species including lipooligosaccharides and cardiolipins, both of which we find to be depleted upon compound treatment. These results are consistent with previous findings regarding SH inhibitor mechanisms. Recent results using a microscopy-based morphological analysis, MorphEUS (Li et al., 2021), concluded that the SH inhibitors AA692 and THL are likely to interfere with cell wall synthesis. In contrast, a different study found that the beta lactone EZ120 caused a decrease the synthesis of mycolic acids, presumably through its inhibition of Ag85 (Lehmann et al., 2018). In our data, Ag85 is competed by JCP276 but surprisingly not by the structurally similar JCP275 and Ag85 is not enriched by BMB034 (Table S1), suggesting that Ag85 may not be the most important target for growth inhibition by the chloroisocoumarins. In further support of this, we did not observe a decrease in mycolic acids upon treatment with JCP276 (Fig. S3c, Table S2).

Although TesA is nonessential for growth in culture, it remains an intriguing target because of its importance for pathogenesis and antibiotic susceptibility. TesA is a critical virulence factor due to its necessity for the synthesis of PDIM. In addition, *tesA* mutants show increased susceptibility to antibiotics (Bellerose et al., 2020; Chavadi et al., 2011; Yang et al., 2020), presumably due to the increased permeability of PDIM-deficient cells. The fact that our wild-type strain contains a mutation in *ppsA* is serendipitous because it allows us to decouple the effects of TesA and other SH inhibition from the general increased sensitivity due to PDIM deficiency. In the case of JCP276 activity, it shows that growth inhibition is not related to the role TesA plays in synthesizing PDIM. However, thioesterase activity is important for the synthesis of other lipids as well. While TesA has been characterized for its role in releasing fatty acids from the PpsABCDE and Mas PKS system required for PDIM synthesis, it is not known whether it interacts with other PKS systems. In addition, while the targets of JCP276 are mostly uncharacterized, it has been shown that the lipase LipG also has thioesterase activity (Santucci et al., 2018).

This study highlights the value of utilizing phenotypic, target-agnostic, screens to identify new pathways that are susceptible to chemical inhibition for the treatment of *M. tuberculosis*. Our lipidomic data and meta-analyses of prior SH inhibitors point to a polypharmacological mechanism of action in which simultaneous inhibition of key SHs prevent bacterial growth. Future screening efforts would benefit from exploring lipid perturbations in addition to growth inhibition as phenotypic outputs. The diversity of SH inhibitors with anti-mycobacterial activity and the evidence that targeting multiple SHs may prevent the evolution of resistance suggests that SHs are potentially important anti-biotic targets for *M. tuberculosis*. Furthermore, future efforts in identifying anti-mycobacterial compounds would likely benefit from targeting a critical set of enzymes that have been prioritized through phenotypic screens and functional studies of their roles in lipid metabolism.

## Acknowledgements

We thank Renier Van der Westhuyzen (Drug Discovery and Development Center, University of Cape Town, South Africa) for assistance with the original phenotypic screen. Mycobacterial clinical isolates were kindly provided by Dr. Nikki Parrish and Derek Armstrong (Department of Pathology, the Johns Hopkins University School of Medicine). We thank Carolyn Bertozzi (Stanford University) for use of the BSL3 facility and mass spectrometry instruments. We thank Nick Riley (Stanford University) and Marc Driessen (Stanford University) for help with mass spectrometry. We thank Jessica Seeliger (Stony Brook University) for helpful discussions and critical feedback. B.M.B. was supported by the A. P. Giannini Foundation. This work was also funded in part by grants from the National Institutes of Health (T32AI07328 to B.M.B., R01 EB005011 to M.B. and R01EB026332 to M.B.). The content is solely the responsibility of the authors and does not necessarily represent the official views of the National Institutes of Health.

## Author Contributions

**Brett M. Babin:** Conceptualization, Data curation, Formal analysis, Investigation, Software, Visualization, Writing – original draft.

**Laura J. Keller:** Data curation, Formal analysis, Software

**Yishay Pinto:** Formal analysis

**Veronica L. Li:** Investigation

**Andrew Eneim:** Investigation

**Summer E. Vance:** Investigation

**Stephanie M. Terrell:** Investigation

**Ami Bhatt:** Funding acquisition, Resources, Supervision

**Jonathan Z. Long:** Funding acquisition, Resources

**Matthew Bogyo:** Conceptualization, Funding acquisition, Resources, Supervision, Writing – review & editing

## Methods

### Bacterial growth conditions

Unless otherwise noted, *M. tuberculosis* was cultured in 7H9/OADC/glycerol medium (4.7 g/l 7H9 powder, 100 ml/l OADC supplement, 4 ml/l 50% (v/v) glycerol, and 2.5 ml/l 20% (v/v) TWEEN 80). OADC was made fresh before adding to the medium (0.5 g/l oleic acid, 50 g/l Albumin fraction V, 20 g/l dextrose, 40 mg/l catalase, and 8.5 g/l NaCl). Cultures were inoculated from 1 ml frozen bacterial stocks and incubated in 1 l roller bottles with rolling at 37°C for 1-2 weeks. *M. tuberculosis* solid cultures were grown on 7H10 medium (19 g/l 7H10 powder, 20 ml 50% (v/v) glycerol, 100 ml/l OADC supplement).

*M. tuberculosis* H37Rv, *Mycobacterium marinum* M, and *Mycobacterium smegmatis* mc^2^155 were a gift from Carolyn Bertozzi (Stanford University). *M. tuberculosis* mc^2^6020 was a gift from Niaz Banaei (Stanford University). The following clinical isolates of NTMs were a gift from Dr. Nikki Parrish and Derek Armstrong (Department of Pathology, the Johns Hopkins University School of Medicine): *Mycobacterium kansasii*, *Mycobacterium intracellulare*, *Mycobacterium scrofulaceum*, *Mycobacterium avium*, and *Mycobacterium abscessus*.

### High-throughput screening, hit validation, and EC_50_ determination

The original screen was performed in 7H9/casitone/glucose medium (4.7 g/l 7H9 powder, 0.3 g/l casitone, 4 g/l glucose, 0.81 g/l NaCl, and 0.05% (v/v) tyloxapol). *M. tuberculosis* H37Rv was grown to OD_600_ of 0.6-0.7, diluted 500-fold into fresh medium, and 50 µl of the diluted culture were added to each well of 96 well-plates containing 50 µl of medium and test compound (final volume: 100 µl, final compound concentration: 20 µM, final DMSO concentration: 1% (v/v)). Each plate contained 8 wells of DMSO as a maximum growth control and 8 wells of 0.15 µM rifampicin as a minimum growth control. Plates were incubated without shaking for 6 d at 37 °C, 10 µl alamarBlue was added to each well, the plate was incubated for an addition 1 d at 37 °C, and viability in each well was scored visually. Hits were validated by treating with ten-point dose response and estimating EC_50_ values. Growth inhibition was tested as above except cultures were grown in 7H9/OADC/glycerol medium and CellTiterBlue (Promega) was used as a measure of cell viability. Viability was quantified by fluorescent measurement (Ex. 560 nm, Em. 590 nm) in a SpectraMax M3 microplate reader (Molecular Devices). For validation experiments, each compound was tested in a single replicate. For JCP276, JCP275, BMB034, and BMB038, each compound was tested in biological triplicate. Fluorescence values were normalized to DMSO (100% viability) and 150 nM rifampicin (0% viability). Data were fit to a two-parameter logistic model to calculate EC_50_ values.

### Growth inhibition measurements

*M. tuberculosis* H37Rv was diluted to OD_600_ of 0.001 and 5 ml of culture was aliquoted into ink well bottles. Cultures were treated with DMSO, 25 µM JCP276 on day 0, or 25 µM on day 0 and an additional 25 µM on day 14. Cultures were grown with shaking at 37 °C for up to 6 weeks. Growth was monitored by transferring cultures to a 96-well plate and measuring OD_600_ in a SpectraMax M3 microplate reader (Molecular Devices). The experiment was performed in biological triplicate.

### MIC measurements

M. tuberculosis strains H37Rv and CDC1551, M. marinum strain M, M. smegmatis strain mc^2^155, and clinical isolates of M. kansasii, M. scrofulaceum, M. intracellulare, M. abscessus, and M. avium were cultured in 7H9/OADC/glycerol medium. M. tuberculosis mc^2^6020 was cultured in 7H9/OADC/glycerol medium supplemented with pantothenate (24 mg/l), L-lysine (80 mg/l), and 0.2 % (w/v) casamino acids. Escherichia coli strain MG1655 and Staphylococcus aureus strain USA300 were cultured in Mueller-Hinton II broth. M. marinum was cultured at 33 °C and all other bacteria were cultured at 37 °C.

Cultures were diluted to OD_600_ 0.001 and 100 µl of each diluted culture was added to 96 well-plates containing JCP276 at various doses (100 nM to 100 µM) and DMSO control wells. Plates were incubated and scored visually for dense bacterial growth. *M. tuberculosis* strains were scored after 3 weeks. All other bacteria were scored after 1 d to 2 weeks after dense growth was observed in the DMSO control wells. MICs were determined as the lowest dose that prevented bacterial growth in all three biological replicates.

### Mammalian cell cytotoxicity measurements

J774 murine macrophages and HFF-1 human foreskin fibroblasts were cultured in DMEM with 10% FBS at 37 °C with 5% CO_2_. Cells were seeded into 96-well plates (20,000 cells/well) and incubated for 1 d. Cells were washed with PBS and resuspended in serum-free DMEM containing DMSO or various concentrations of JCP276 (1% maximum DMSO concentration). Plates were incubated for 24 h and viability was measured following a 6 h incubation with CellTiterBlue (Promega). Viability was quantified by fluorescent measurement (Ex. 560 nm, Em. 590 nm) in a Cytation 3 microplate reader (Biotek). Fluorescence values were normalized to DMSO (100% viability) and wells containing no cells (0% viability). The experiment was performed in biological triplicate.

### ABPP labeling and enrichment

Cultures of *M. tuberculosis* H37Rv were centrifuged and resuspended to a final OD_600_ of 2.5 in 7H9 buffer (4.7 g/l). Cultures were treated with JCP275, or JCP276, or DCI and incubated at 37°C for 1 h. For FP labeling, cells were lysed (see below) and lysates were treated with 1 µM FP-TMR or 5 µM FP-biotin for 30 min at 37 °C. For BMB034 labeling, cells were treated with 1 µM BMB034 for 1 h at 37 °C immediately after compound treatment and then lysed.

Bacteria were resuspended in PBS and lysed by bead beating at room temperature. Triton-X was added to a final concentration of 1% and lysates were clarified by centrifugation at 2800 rcf for 5 min at room temperature. Lysates were filtered twice through 0.2 µm syringe filters for removal from the BSL3 facility.

For FP-TMR experiments, lysates were analyzed directly by SDS-PAGE. For BMB034 experiments, lysates were subjected to click chemistry conditions (5 mM BTTAA, 1mM CuSO_4_, and 15 mM sodium ascorbate) with 20 µM azide-TMR for 30 min at 37 °C and analyzed by SDS-PAGE. For BMB034 enrichments, lysates were subjected to click chemistry conditions with 20 µM desthiobiotion-TMR-azide for 1 h at 37 °C.

Biotinylated protein samples were desalted using PD-10 columns. Proteins were eluted into PBS and SDS was added to a final concentration of 0.05%. Samples were heated at 90 °C for 8 min and then cooled at -20 °C for 5 min. Streptavidin-agarose was washed in PBS and 100 µl of slurry was added to each protein sample and allowed to mix for 1 h at room temperature. Agarose resin was pelleted by centrifugation at 4000 rcf for 5 min between each of the following washes: twice with 1 % SDS, twice with 6 M urea, and twice with PBS. Beads were aspirated and then resuspended in 500 µl 6 M urea. Proteins were reduced by addition of 25 µl of 30 mg/ml DTT and incubation for 15 min at 65 °C. Samples were cooled and proteins were alkylated by addition of 25 µl of 14 mg/ml iodoacetamide and incubation for 30 min at 37 °C. Beads were rinsed once with 1 ml PBS and resuspended in digestion buffer (200 µl 2 M urea, 2 µl 1 M CaCl_2_, 4 µl of 0.5 mg/ml trypsin). Proteins were digested for 16 h at 37 °C.

Beads were transferred to spin filters and washed twice with 50 µl PBS to yield digested peptides. Peptides were acidified by addition of 15 µl formic acid and desalted using C18 zip tips. Peptides were eluted into 75% acetonitrile with 1% formic acid, dried, and stored at -20 °C for subsequent LC-MS/MS analysis.

### Proteomic analysis

Peptides were resuspended in 0.2% formic acid in water. Peptides were separated over a 25- cm EasySpray reversed phase LC column (75 μ m, 100 Å, PepMap C18 particles, Thermo Fisher Scientific). The mobile phases (A: water with 0.2% formic acid and B: acetonitrile with 0.2% formic acid) were driven and controlled by a Dionex Ultimate 3000 RPLC nano system (Thermo Fisher Scientific). Gradient elution was performed at 300 nl min^−1^. Mobile phase B was increased to 5% over 6 min, followed by an increase to 40% at 60 min, a ramp to 90% B at 61 min, and a wash at 90% B for 10 min. Eluted peptides were analyzed on an Orbitrap Fusion Tribrid MS system (Thermo Fisher Scientific). Precursors were ionized using an EASY-Spray ionization source (Thermo Fisher Scientific) source held at +2.2 kV compared to ground, and the column was held at 40°C. The inlet capillary temperature was held at 275°C. Survey scans of peptide precursors were collected in the Orbitrap from 350-1500 Th with an automatic gain control target of 1,000,000, a maximum injection time of 50 ms and a resolution of 120,000 at 200 m/z. Monoisotopic precursor selection was enabled for peptide isotopic distributions, precursors of z = 2–5 were selected for data-dependent MS/MS scans for 2 s of cycle time, and dynamic exclusion was set to 45 s with a ±10 ppm window set around the precursor monoisotope. An isolation window of 0.7 m/z was used to select precursor ions with the quadrupole. MS/MS scans were collected using high-energy collisional dissociation at 30 normalized collision energy (nce) with an automatic gain control target of 30,000 and a maximum injection time of 25 ms. Mass analysis was performed in the linear ion trap using the ‘Rapid’ scan speed while scanning from 200–1,500 m/z.

### Proteomic data analysis

Raw data files were searched and quantified using MaxQuant LFQ. Data were searched against the reference H37Rv proteome. FP and BMB034 enrichment experiments were analyzed separately and results were merged (Table S1). Protein abundance ratios were calculated by dividing LFQ values from compound-treated samples by those from DMSO-treated samples. In cases where proteins were not identified in compound-treated samples, log_2_ ratios were arbitrarily set to -10.

### Lipid extractions

*M. tuberculosis* H37Rv liquid cultures at OD_600_ of 1 were transferred to 0.22 µm membranes placed on 7H10 solid medium and incubated for 1 week at 37 °C to yield a lawn of bacteria on each membrane. Membranes were transferred to 6-well plates containing 7H10 solid medium with 1% DMSO, 100 µM JCP276, 100 µM BMB034, or 100 µM THL. Membranes were incubated for 24 h at 37 °C. Each membrane was transferred to an O-ring tube and lipids were extracted form whole cells by vortexing once each with 0.5 ml 1:2 chloroform:methanol, 0.5 ml 1:1 chloroform:methanol, and 0.5 ml 2:1 chloroform:methanol. Supernatants from the three extractions were combined and filtered twice through 0.22 µm syringe filters before removal from the BSL3 facility. Extracts were transferred to glass vials and solvent was dried under nitrogen gas. Samples were dried completely by rotary evaporation. The dry mass of each lipid sample was measured and lipids were resuspended in 2:1 chloroform:methanol to a final concentration of 5 mg/ml.

### Lipidomic analysis

Mass spectrometry analysis was performed with an electrospray ionization source on an Agilent 6545 Q-TOF LC/MS in negative ionization mode. For Q-TOF acquisition parameters, the mass range was set from 100 to 3000 m/z with an acquisition rate of 10 spectra/second and time of 100 ms/spectrum. For Dual AJS ESI source parameters, the drying gas temperature was set to 250°C with a flow rate of 12 l/min, and the nebulizer pressure was 20 psi. The sheath gas temperature was set to 300°C with a flow rate of 12 l/min. The capillary voltage was set to 3500 V and the fragmentor voltage was set to 100 V. Reversed-phase chromatography was performed with a Luna 5 mm C5 100 Å LC column (Phenomenex cat #00B-4043-E0). Samples were injected at 20 ul each. Mobile phases for were as follows: Buffer A, 95:5 water/methanol with 0.1% ammonium hydroxide; Buffer B, 60:35:5 isopropanol/methanol/water with 0.1% ammonium hydroxide. All solvents were HPLC-grade. The flow rate for each run started with 0.5 minutes 95% A / 5% B at 0.6 ml/min, followed by a gradient starting at 95% A / 5% B changing linearly to 5% A / 95% B at 0.6 ml/min over the course of 19.5 minutes, followed by a hold at 5% A / 95% B at 0.6 ml/min for 8 minutes and a final 2 minute at 95% A / 5% B at 0.6 ml/min.

### Lipidomic data analysis

Raw files were converted to mzXML format with MSConvert (ProteoWizard) using the Peak Picking Vendor algorithm. Files were analyzed using the web-based XCMS platform (Tautenhahn et al., 2012) with the following settings: signal to noise threshold, 6; maximum tolerated m/z deviation, 30 ppm; frame width for overlapping peaks, 0.01; and peak width, 10-60 s. Integrated peak intensities were normalized between conditions by median fold change.

Tables containing mass features and ion intensities from each experiment were downloaded from the XCMS platform. Identified ions were matched to the MycoMass database (Layre et al., 2011) using a custom python script. Ions were matched if the measured m/z was within 10 ppm of the annotated m/z.

### Generation of resistant mutants

*M. tuberculosis* H37Rv was grown to an OD_600_ of 1, centrifuged, and concentrated 10-fold (∼10^9^ CFU/ml). Concentrated cultures (1 ml) were added to 7H10 solid medium containing 100 µM JCP276 and allowed to incubate for 3 weeks. Bacteria grown in the presence of JCP276 from three separate plates were used to start liquid cultures which were then tested for JCP276 susceptibility and frozen for future use (annotated BBMT04, BBMT05, and BBMT06).

### Whole genome and sanger sequencing of mutants

Cultures of H37Rv, BBMT04, BBMT05, and BBMT06 were grown in liquid culture to an OD_600_ of 1. Cultures were centrifuged and resuspended in breaking buffer (50 mM Tris pH 8, 10 mM EDTA, and 100 mM NaCl). Bacteria were lysed by bead beating, the supernatant was transferred to a clean O-ring tube, and the samples were adjusted to 1% final concentration of SDS. Samples were filtered through 0.22 µm syringe filters twice and removed from the BSL3 facility. DNA was purified using a Qiagen cleanup kit. The *ppsA* region of interest was amplified by PCR using primers *ppsA-seq.for* (TCGACGCGGAATTCTTCGAG) and *ppsA-seq.rev* (AACCGCTTGAGCACCACTAC) and analyzed by Sanger sequencing.

### Purification and FP competition of recombinant TesA

The *tesA* gene was amplified by PCR from chromosomal H37Rv DNA and cloned into pET28b to generate pET28b-tesA, encoding a 6xHis tag on the N-terminus of the protein. The expression vector was transformed into *E. coli* BL21/DE3. Expression cultures were grown in 2x YT medium to an OD_600_ of 0.4, induced with 1 mM IPTG, and allowed to express for 16 h at 20 °C. Cultures were centrifuged and the combined pellet was lysed by sonication in lysis buffer (50 mM NaH_2_PO_4_ pH 7.5, 300 mM NaCl, 5 mM imidazole). The lysate was clarified by centrifugation and the soluble protein was purified by a HisTrap column on an Akta chromatography system, using gradient elution from 100% lysis buffer to 100% elution buffer (50 mM NaH_2_PO_4_ pH 7.5, 300 mM NaCl, 300 mM imidazole). Fractions containing TesA were combined and further purified by gel filtration with a Superdex 75 column using GF buffer (50 mM NaH_2_PO_4_ pH 7.5, 100 mM NaCl).

Purified TesA (100 nM) was incubated with DMSO or various concentrations of JCP276 for 30 min at 4 °C and was then treated with 1 µM FP-TMR for 15 min at 4 °C. The sample was immediately quenched by addition of SDS buffer and separated by SDS-PAGE.

### Software

Plotting and statistical analysis were performed with Prism (GraphPad) or python with numpy, scipy, matplotlib, pyplot, and seaborn packages. Proteomics data were analyzed using MaxQuant. Lipidomics data were analyzed using msConvert and the XCMS online platform.

## Supplemental Tables

**Table S1. Proteins identified in ABPP experiments.** Proteins are indicated by their UNIPROT ID, locus ID, protein name and description, and predicted molecular weight (MW). MaxQuant LFQ values are given for each experimental condition (average of three biological replicates). Log_2_(LFQ ratios) were calculated by dividing the LFQ value for each compound-treated sample by the LFQ value for the DMSO-treated controls. For proteins that were not detected in the compound-treated sample, the log_2_(ratio) was set to -10. Proteins identified by FP pulldown in this study were compared to those identified in prior ABPP experiments (Li et al., 2021; Ortega et al., 2016; Tallman et al., 2016).

**Table S2. Lipids identified in LC-MS lipidomic experiments.** Lipid species are indicated by their unique Ion identifiers, retention time, and m/z. Number of matches per replicate and mean ion intensities are indicated for each experimental condition (average of three biological replicates). Ions were matched to lipid annotations in the MycoMass database (Layre et al., 2011) yielding class, subclass, abbreviation, chemical formula, and molecular weight. The matched ion is indicated. Log_2_(intensity ratios) were calculated by dividing the mean ion intensities for each compound-treated sample by the mean ion intensity for the DMSO-treated controls. For lipids that were only detected in the DMSO or compound-treated samples, the log_2_(intensity ratio) was set to -10 or 10, respectively.

**Table S3. Mutations identified in the wild-type strain.** Each line represents a mutation identified in the wild-type strain as compared to the H37Rv reference genome (NCBI RefSeq: NC_000962.3). The base(s) at each position in the reference genome (“Reference sequence”) are replaced by the bases confirmed by sequencing (“Confirmed sequence”). When the mutation occurs within a coding region (i.e., not intergenic), the locus and gene name are indicated and the effect of the mutation on the coding sequence (“Consequence”) is noted: *, stop codon; fs, frame shift; del, amino acid deletion; orf, open reading frame.

