## Supplementary Figures for "A screen of covalent inhibitors in *Mycobacterium tuberculosis* identifies serine hydrolases involved in lipid metabolism as potential therapeutic targets"

**Supplemental Figures**
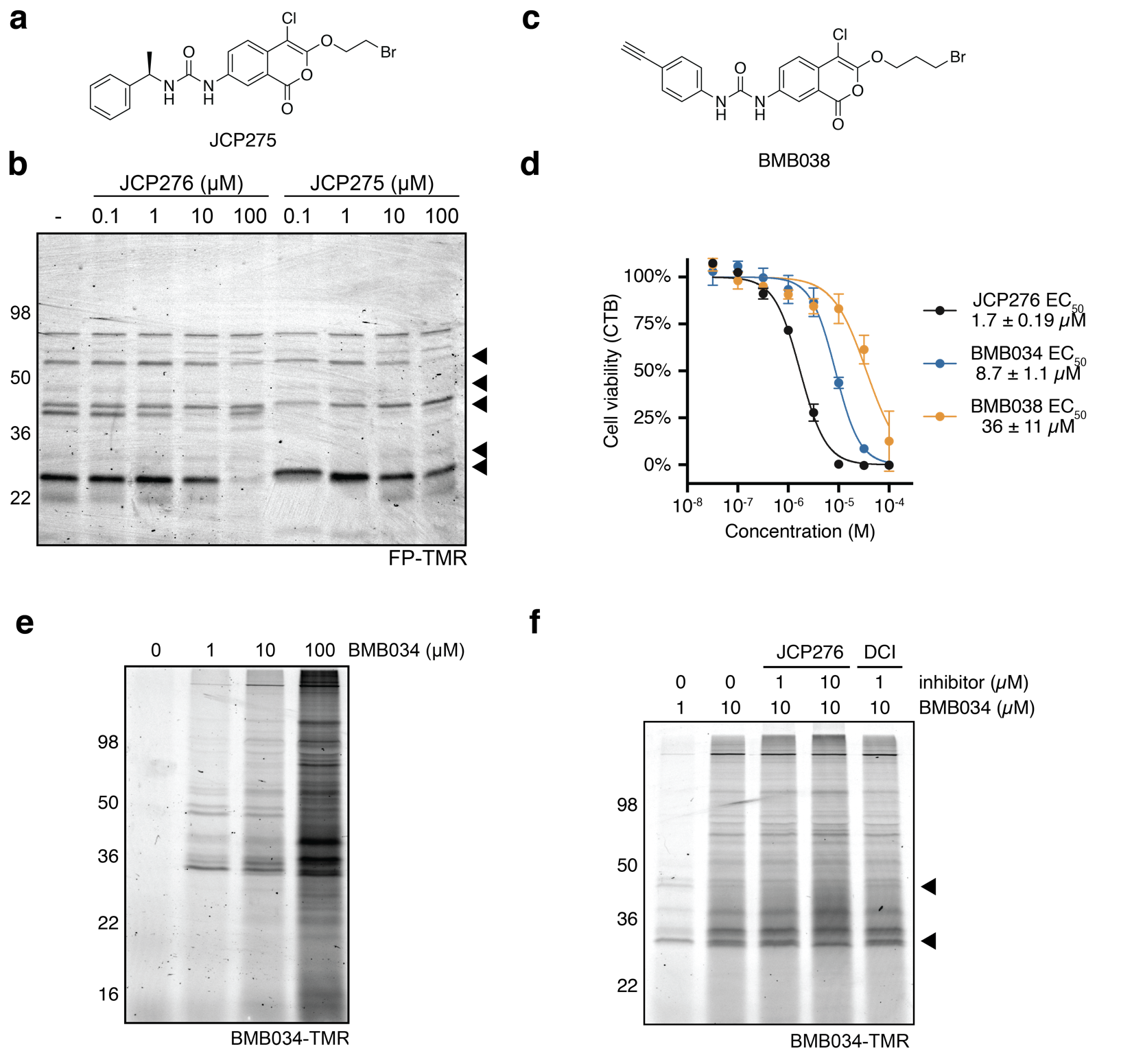

**Figure S1. FP and BMB034 labeling of *M. tuberculosis* lysates.** (a) Structure of JCP275. (b) Dose-dependent competition of FP-TMR labeling by JCP276 and JCP275. Bacterial cultures were treated with DMSO or JCP276 as indicated for 1 h at 37 ºC. Lysates were treated with 1 µM FP-TMR for 30 min at 37 ºC and proteins were separated by SDS-PAGE. (c) Structure of BMB038. (d) Dose-dependent inhibition of *M. tuberculosis* growth by JCP276 following a 7-day incubation. Data represent the mean ± standard deviation of three biological replicates. Data were fit to two-parameter parametric model (solid line) and the EC_50_ is reported as the mean ± 95% confidence interval. (e-f) Competitive ABPP using BMB034. Bacterial cultures were pre-treated with DMSO, JCP276, or DCI as indicated for 1 h at 37 ºC and then treated with BMB034 as indicated for 1 h at 37 ºC. Lysates were subjected to click chemistry conditions with azide-TMR and proteins were separated by SDS-PAGE. Gels show TMR fluorescence.

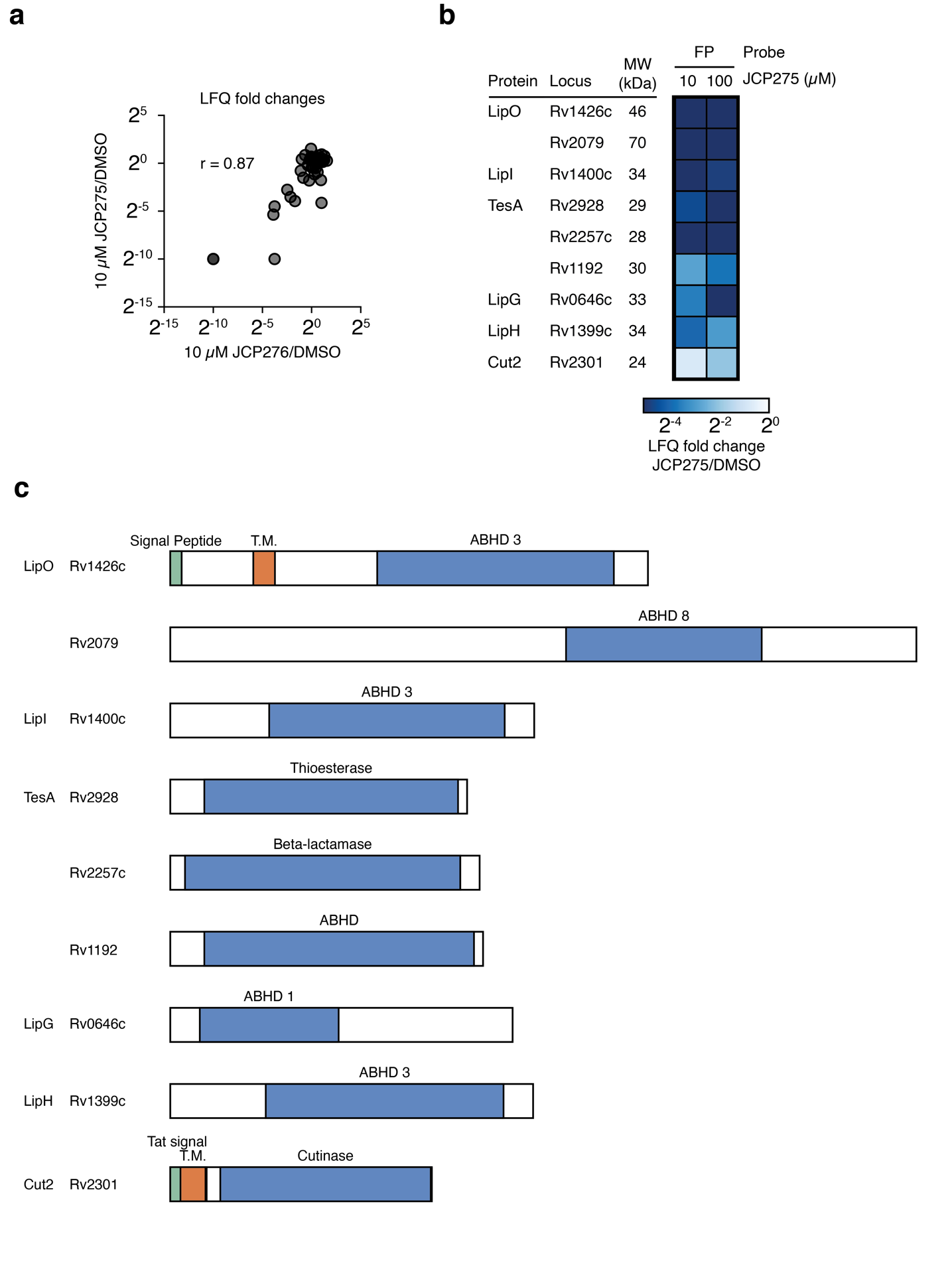

**Figure S2. ABPP proteomic results.** (a) A comparison of LFQ ratios for proteins enriched by FP-biotin following pre-treatment with 10 µM JCP275 or 10 µM JCP276. The Pearson correlation coefficient (r) is indicated. (b) Summary of proteomic results from the FP ABPP experiment with JCP275. The heatmap represents the ratio of each protein’s abundance between cultures treated with JCP275 at the indicated dose and cultures treated with DMSO. (c) Domain annotations for each protein target. SH domains are shown in blue.

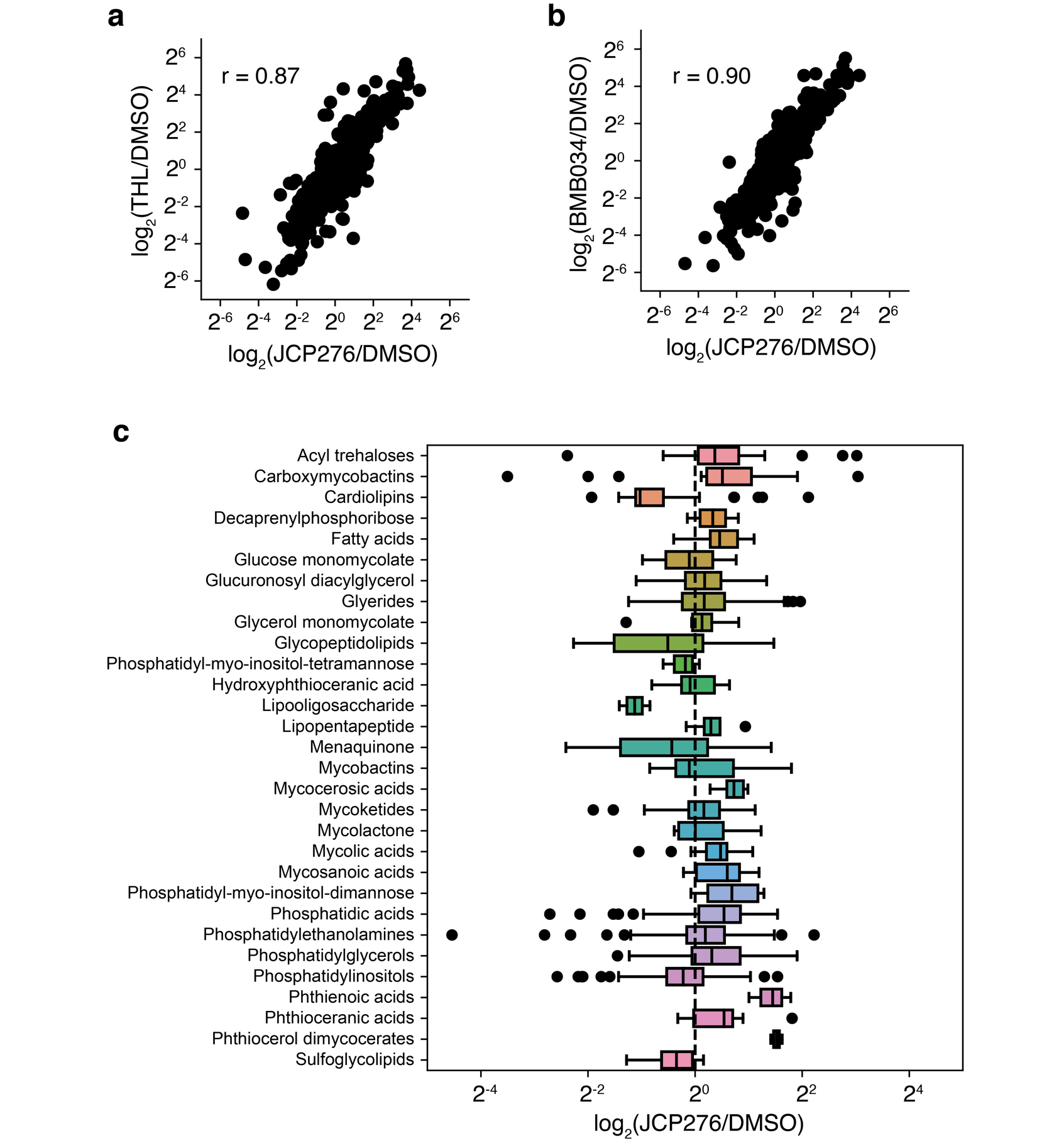

**Figure S3. Lipidomic results.** (a, b) Comparisons of ion intensity ratios between compound-treated and DMSO-treated cultures for (a) THL vs. JCP276 and (b) BMB034 vs. JCP276.The Pearson correlation coefficients (r) are indicated. (c) Boxplots of lipid abundance ratios between cultures treated with each compound or cultures treated with DMSO. The vertical line represents the median value for each lipid class, boxes represent the upper and lower quartiles, whiskers represent 1.5 times the interquartile range, and circles represent outliers from this range.
